# The effect of memantine, an antagonist of the NMDA glutamate receptor, in *in vitro* and *in vivo* infections by *Trypanosoma cruzi*

**DOI:** 10.1101/546861

**Authors:** Higo Fernando Santos Souza, Sandra Carla Rocha, Flávia Silva Damasceno, Ludmila Nakamura Rapado, Elisabeth Mieko Furusho Pral, Claudio Romero Farias Marinho, Ariel Mariano Silber

## Abstract

Chagas disease, caused by *Trypanosoma cruzi,* is a neglected tropical disease that affects 5-6 million people in endemic areas of the Americas. Presently, chemotherapy relies on two compounds that were proposed as trypanocidal drugs four decades ago: nifurtimox and benznidazole. Both drugs are able to eliminate parasitemia and to avoid seroconversion in infected people when used in the acute phase; however, their use in the chronic phase (the time when the majority of cases are diagnosed) is limited due to their serious side effects. Memantine is a glutamate receptor antagonist in the central nervous system of mammals that has been used for the treatment of Alzheimer’s disease. Our group previously reported memantine as a trypanocidal drug that is able to induce apoptosis-like death in *T. cruzi*. In the present work, we further investigated the effects of memantine on the infection of RAW 264.7 macrophages *in vivo* (in BALB/c mice). Here, we showed that memantine is able to diminish NO and Ca^2+^ entry in both LPS-activated and non-activated cells. These results, together with the fact that memantine was also able to reduce the infection of macrophages, led us to propose that this drug is able to activate a pro-oxidant non-NO-dependent cell defense mechanism. Finally, infected mice that were treated with memantine had diminished parasitemia, cardiac parasitic load, and inflammatory infiltrates. In addition, the treated mice had an increased survival rate. Taken together, these results indicate memantine to be a candidate drug for the treatment of Chagas disease.

**Author summary:** Chagas disease affects approximately 5 million people and is caused by the protist parasite *Trypanosoma cruzi.* Until now, there are no vaccines to prevent the human infection, and the therapy relies on the use of two drugs discovered more than 50 years ago, nifurtimox and benznidazole. Both drugs are efficient during the acute phase of the disease, however their efficacy in the chronic phase, when most of patients are diagnosed is controversial. In addition, both drugs are toxic, causing severe side effects during the treatment. For these reasons, new drugs against *T. cruzi* are urgently needed. In this work, we report a series of experiments supporting the repositioning of memantine, a drug used for treating Alzheimer’s disease, to treat the *T. cruzi* infection in an experimental infection model. Our data show that infected mice treated with memantine have diminished their parasitemia, cardiac parasitic load and inflammatory infiltrates and more importantly, they have diminished their mortality. Taken together, these results prompt memantine as a promising drug for treating Chagas disease.

## Introduction

Chagas disease is caused by the protozoan *Trypanosoma cruzi* and affects 5-6 million people in the Americas (1). Mammals (including humans) become infected when an infected triatomine insect defecates on the skin and expels metacyclic trypomastigotes with the feces, one of the nonproliferative, infective forms of the parasite. These forms are able to internalize into mammalian hosts through the mucosa and small wounds caused by scratching. Once inside the mammalian host, the metacyclic trypomastigotes invade the host cells to reach the cytoplasm, where they initiate their proliferation as amastigotes. After a variable number of cellular divisions, amastigotes undergo a complex differentiation process, yielding a new generation of infective, nonproliferative forms called trypomastigotes. These trypomastigotes burst from the infected cells into the extracellular environment and are able to infect the neighboring cells or to reach the bloodstream, allowing them to extend the infection to other tissues. Eventually, the bloodstream trypomastigotes can be taken by the blood to a new, noninfected triatomine insect during its blood-meal, can infect the insect, and can convert this newly infected insect into a new transmitter of the infection (2).

Chagas disease can be divided into two phases: acute and chronic. The acute phase is mainly asymptomatic with evident parasitemia and undetectable levels of IgG antibodies. The chronic phase is characterized by a robust humoral response with high titers of IgG antibodies and subpatent parasitemia. The chronic phase persists for the host’s lifespan. Most patients in the chronic phase (60-70%) are asymptomatic. However, the remaining 30-40% of chronic patients develop recognizable clinical symptoms. The most frequent symptoms are heart hypertrophy and dilatation, esophagus and large intestine dilatations (megavisceras), or a combination of both (reviewed by (3, 4)). The treatment of the chagasic infection is largely unsatisfactory (5). Presently, two drugs discovered approximately 50 years ago are available nifurtimox (Nf) and benznidazole (Bz). Both drugs are highly effective in the acute phase. However, their efficacy in treating the chronic phase, when most patients are diagnosed, is limited due to the serious side effects that occur from the toxicity of the drugs and the long-term treatment required in this phase. Importantly, the emergence of resistant parasites was reported. In view of these facts, there is an urgent need to look for new drugs to treat *T. cruzi* infections (3, 6).

Our group has been exploring drug reposition strategies, consisting of the identification of new uses for drugs already approved for the treatment of any disease in humans (7, 8). In a previous work, Paveto et al. suggested the existence of an L-glutamate receptor N-methyl-D-aspartate (NMDA) type in *T. cruzi*, which would be analogous to those reported in neural cells (9). Additionally, our group characterized a *T. cruzi* glutamate transporter (10) that could behave as a glutamate receptor. More recently, we showed the sensitivity of *T. cruzi* to memantine (1,2,3,5,6,7-hexahydro-1,5:3,7-dimethano-4-benzoxonin-3-yl) amines, a tricyclic amine with a low-to-moderate affinity for the *N*-methyl-D-aspartate (NMDA) receptor (11), which has been indicated for the treatment of Alzheimer’s disease (12). More specifically, we showed that memantine presented an apoptotic-like activity in *T. cruzi* epimastigotes as well as a trypanocidal effect in infected mammalian cells (11). In the present work, we show that memantine affects the infection of macrophages by *T. cruzi,* diminishing the number of infected cells. We also report that, in addition to its effect on the parasite, memantine modifies macrophage activation by slightly diminishing both NO production and intracellular Ca^2+^ levels in activated and non-activated macrophages. Finally, infected mice treated with memantine presented a diminished parasitemia peak, heart parasitic load, inflammatory infiltrates, and mortality. As a whole, this work proposes memantine to be an interesting drug to be further explored for the treatment of Chagas disease.

## Materials and Methods

### Reagents

Memantine was purchased from Tocris Bioscience (Minneapolis, MN, USA). The DNA extraction kit, DNAeasy Blood and Tissue Kit, was purchased from Qiagen (Hilden, DE). Culture medium and fetal calf serum (FCS) were purchased from Cultilab (Campinas, SP, Brazil). Fluo-4 AM were purchased from Invitrogen (Eugene, Oregon, USA). The MTT [3-(4,5-dimethylthiazol-2-yl)-2,5-diphenyltetrazolium bromide] assay, the bioluminescent somatic cells kit, lipopolysaccharide from *Escherichia coli* (LPS) and Griess reagent were purchased from Sigma-Aldrich (St. Louis, MO, USA). The dichloro-dihydro-fluorescein diacetate (DCFH-DA) assay, Reverse Transcription *SuperScriptII* kit, Trizol reagent, SYBR Green Master Mix, fluo-4 AM and Hoechst 33258 were purchased from Thermo Fisher Scientific (Carlsbad, CA, USA).

### Animals

Six- to eight-week-old BALB/c female mice were obtained from the animal facility of the Department of Parasitology of ICB, USP. The animals were kept under controlled climatic conditions with free access to food and water (*ad libitum*). All laboratory procedures involving animals were previously authorized by the Ethics Committee on Animal Use for ICB-USP (Protocol 107, Fls 132, Book 02).

### Mammalian cells and parasites

The RAW 264.7 (macrophage) cell line was routinely cultivated in RPMI 1640 medium supplemented with 10% heat-inactivated fetal calf serum (FCS), supplemented with 2 mM sodium pyruvate, 0.15% (w/v) NaCO_3_, 100 units mL^-1^ penicillin and 100 µg mL^-1^ streptomycin at 37 °C in a humid atmosphere containing 5% CO_2_. Tissue culture-derived trypomastigotes of the *T. cruzi* Y-strain were obtained from infections of the LLC-MK_2_ cell line (multiplicity of infection: 10 trypomastigotes/cell) as previously described (13). Trypomastigotes were collected from the supernatant of LLC-MK_2_ cells at days 6 to 10 postinfection and were transferred to other bottles for new passages and/or used for infection assays. The bloodstream trypomastigote form of the *T. cruzi* Y-strain was maintained by infecting the BALB/c mice. The recovery of trypomastigotes was performed weekly and was used for the infection assays.

### Determination of RAW 264.7 macrophage viability

RAW 264.7 cells (5.0 x 10^5^ cells mL^-1^) were cultured in 24-well plates in RPMI medium supplemented with FCS (10%) in the presence of different concentrations of memantine (ranging from 10 to 800 μM) or none (control). Cell viability was evaluated 48 h after the initiation of treatment using an MTT assay (3-(4,5-dimethylthiazol-2-yl)-2,5-diphenyltetrazolium bromide) (14). The inhibitory concentration of 50% of cells (IC_50_) was determined by fitting the data to a typical dose-response sigmoidal curve using the program OriginPro8.

### Effect of memantine on the intracellula*r* amastigote of *T. cruzi*

RAW 264.7 cells (2.5 x 10^4^ per well) were cultivated on coverslips in 24-well plates in RPMI medium (10% FCS) and kept at 37 °C. After 24 h, the cells were infected with the trypomastigote form of the Y-strain (2.5 x 10^5^ per well) for 4 h. After this time, free parasites were removed by washing twice with PBS; RPMI medium (10% FCS) was replaced, and the cells were treated with different concentrations of memantine (range 10 µM to 100 µM) for the following 72 h. Then, the cells were washed with PBS, fixed with 4% paraformaldehyde for 5 min, washed again and treated with Hoechst 33258 for 1 min. After washing, the cells were observed by fluorescence microscopy. The infected cells were counted from a sample of 400 randomly chosen cells for determination of the infection rate. The number of amastigotes was also counted to determine the rate of amastigote per cell.

### Evaluation of the nitric oxide (NO) production from RAW 264.7 macrophage culture

Macrophages require activation by particular quantities of *Escherichia coli-*derived LPS for NO production detection. Thus, a dose-response curve was produced to determine the ideal concentration of LPS required for activating the RAW 264.7 cells. The macrophages were stimulated with different concentrations of LPS ranging from 1 to 100 μg/ml for 24, 48 and 72 h. The NO evaluation was based on the nitrite leased measure from the supernatant of the cultured cells. The RAW 264.7 cells (2.5 x 10^5^ cells mL^-1^) were cultured in 96-well plates in RPMI medium (10% FCS) in the presence of different concentrations of memantine (ranging from 1 to 100 μM) or none (control) and were stimulated by 10 μg/ml LPS for 24 h or no stimulation. Over this period, the nitrite concentration of the cell supernatant was quantified using the Griess reaction, as described by (15).

### Evaluation of the gene expression of inducible NO synthase in RAW 264.7 macrophages

To evaluate the gene expression of inducible nitric oxide synthase, 2 x 10^6^ cells per well were cultured for 18 h in 6-well plates in RPMI medium (10% FCS) in the presence of different concentrations of memantine (ranging from 1 to 100 μM) or none (control) and were stimulated by 10 μg/ml LPS for 18 h. After the incubation time, the supernatant was discarded, and the adhered cells were homogenized with Trizol for RNA extraction (Thermo Fisher Scientific). cDNA was synthetized using the Reverse Transcription Kit *SuperScriptII* (Thermo Fisher Scientific). qPCR was performed with *SYBR Green* (Fermentas) for detecting the gene expression levels of iNOS. All reactions were run in triplicate on an Eppendorf RealPlex Real Time PCR System (Eppendorf) with the standard thermal cycling conditions. The runs were normalized with the ACT-β gene. The threshold cycle (2^-ΔΔCt^) method of comparative PCR was used for the data analysis.

### Analysis of intracellular Ca^2+^ levels in RAW 264.7 macrophages

Cells (2.5 x 10^4^ cells/well) were cultivated in 96-well plates in RPMI medium (10% FCS), stimulated or not stimulated (control) with LPS, and treated or not treated (control) with different concentrations of memantine (ranging between 1 and 100 μM) for 24 h. Then, 5 µM fluo-4 AM (Invitrogen) was added to the cultures for 1 h. After the incubation, the cells were washed twice with HEPES-glucose (50 mM HEPES, 116 mM NaCl, 5.4 mM KCl, 0.8 mM MgSO_4_, 5.5 mM glucose and 2 mM CaCl_2_, pH 7.4). The reading was performed on a Spectra Max M3 fluorometer, Molecular Devices, using excitation λ 490 nm and emission λ 518 nm (11).

### Evaluation of parasitemia and survival of BALB/c mice

Blood samples were obtained from the tails of *T. cruzi*-infected mice (1×10^3^ trypomastigotes per mouse) treated with or without memantine. On the days of parasitemia peaks, the number of trypomastigotes was quantified in 25 microscope fields at 400x magnification with 10 μL of blood (Nikon Eclipse E200) (16). The survival of the mice was also monitored for 40 days postinfection.

### Quantification of the tissue parasite load

On the 15^th^ day postinfection, samples of lung, spleen, bladder, heart, intestine, and skeletal muscle were obtained from the infected BALB/c mice for the quantification of the parasite load. The fragments were transferred to formaldehyde (10%) and were then processed by gradual dehydration in ethanol solutions, followed by immersion in xylene, and subsequently embedded in paraffin. Tissue sections 5 μm thick were obtained and stained with hematoxylin and eosin (H&E) and analyzed by light microscopy. The number of amastigote nets was counted in 20 random microscope fields using a 400x magnification. In parallel, tissue fragments were submitted for DNA extraction using the DNAeasy Blood and Tissue Kit as recommended by the manufacturer. The tissue parasitic load was also performed using quantitative PCR as previously described (17). The cycle threshold values obtained by the Eppendorf RealPlex software were converted to the number of parasites per 5 ng of tissue DNA. Their averages were normalized according to the TNF-α gene.

### Histopathological analysis in the cardiac tissue

Cardiac tissue sections 5 μm thick were obtained on the 15^th^ day postinfection, stained with H & E and analyzed by light microscopy. Six nonconsecutive slides from the heart of each mouse were analyzed in a blinded fashion. Areas of inflammatory infiltrates were quantified by an image analysis system (Bioscan Optimas; Bioscan Inc., Edmonds, Wash). The sum of the infiltrated areas from the six slides was calculated for each mouse. The final individual score was expressed in square micrometers of inflammatory infiltrates per square millimeter of area examined.

### Statistical analysis

The experimental data were input into GraphPad Prism version 4.0 software for construction of the graphs. In addition, t-tests or one-way ANOVA analyses were performed, followed by Tukey’s and logrank tests for statistical analysis. Differences with a *p* value <0.05 were considered statistically significant.

## Results

### In vitro

#### Memantine affects the intracellular cycle of *T. cruzi*

To verify the effect of memantine on the intracellular cycle of *T. cruzi*, the effect of the treatment after the infection was evaluated. First, we evaluated the toxicity of memantine in macrophages of the RAW 264.7 lineage. The cells were treated with different concentrations of memantine (10 – 800 µM) for 24, 48 and 72 h; we observed that the macrophages tolerated memantine at concentrations up to 100 μM, showing an IC_50_ of 580 ± 22 µM, 279 ± 2 µM and 257 ± 4.7 µM, respectively (**Fig 1A-F**).

**Figure 1.**
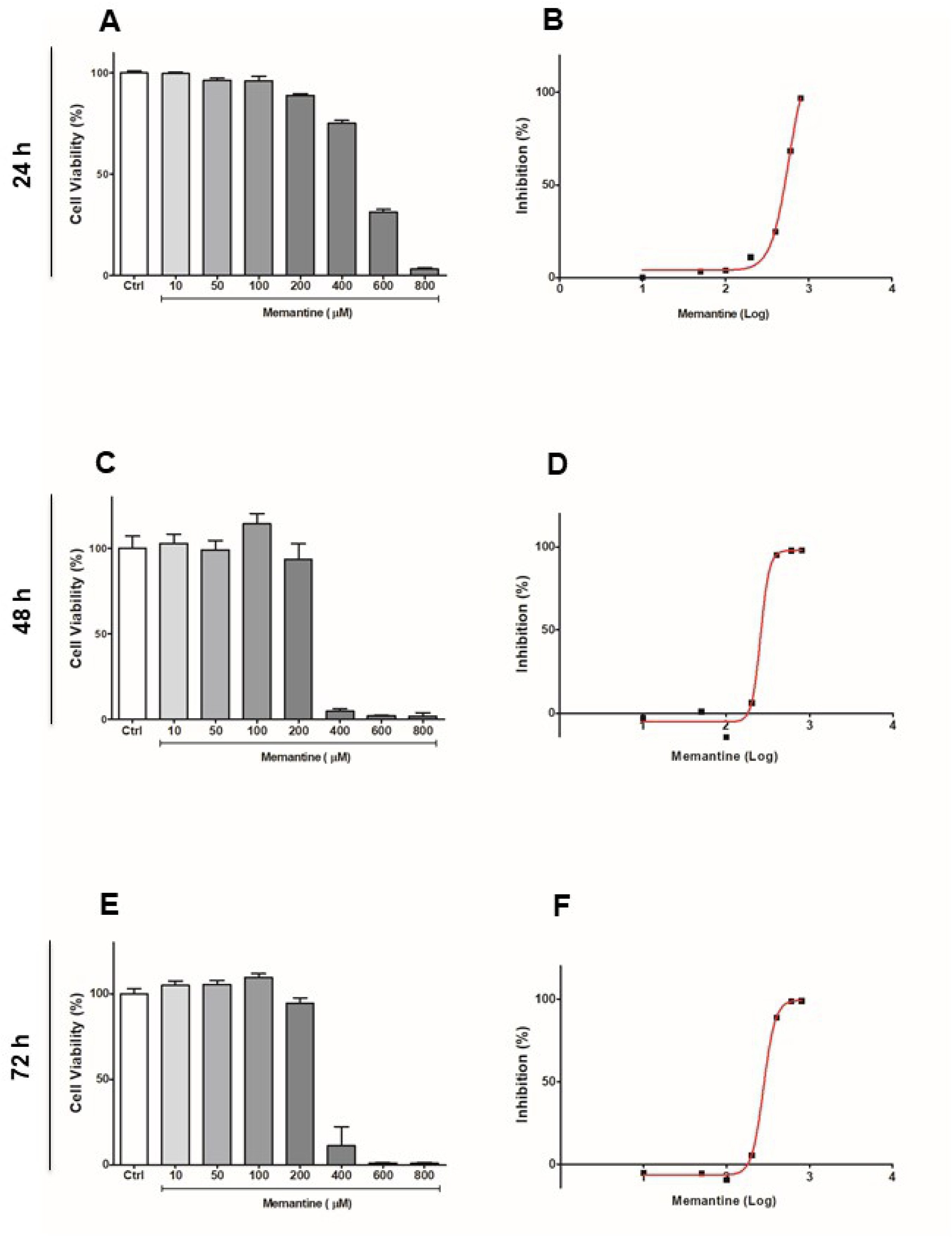
Evaluation of cell viability in RAW 264.7 cells. **(A, C and E)** The cells were treated with different concentrations of memantine (10-800 μM), incubated for 24, 48 and 72 hours. **(B, D and F)** IC_50_ values were obtained from a nonlinear regression curve; the determined IC_50_ values were 580 ± 22 μM, 279 ± 2 μM and 257 ± 4.7 μM, respectively. Comparison among the groups treated and not treated with memantine (p <0.05). Data are expressed as a percentage ± standard deviation.

Once the cytotoxic effect of memantine was evaluated, the RAW 264.7 macrophages were subjugated to infection. For this, the cells were incubated with trypomastigotes for 3 h, and then the trypomastigotes remaining in the supernatants were washed out. The infected cells were incubated for 12 h at 37 °C. Then, the culture medium was replaced with culture medium containing different concentrations of memantine (1-100 μM). These treatments were maintained for 72 h. The treatment reduced the number of infected cells at all concentrations tested when compared to the control group (**Fig 2**).

**Figure 2.**
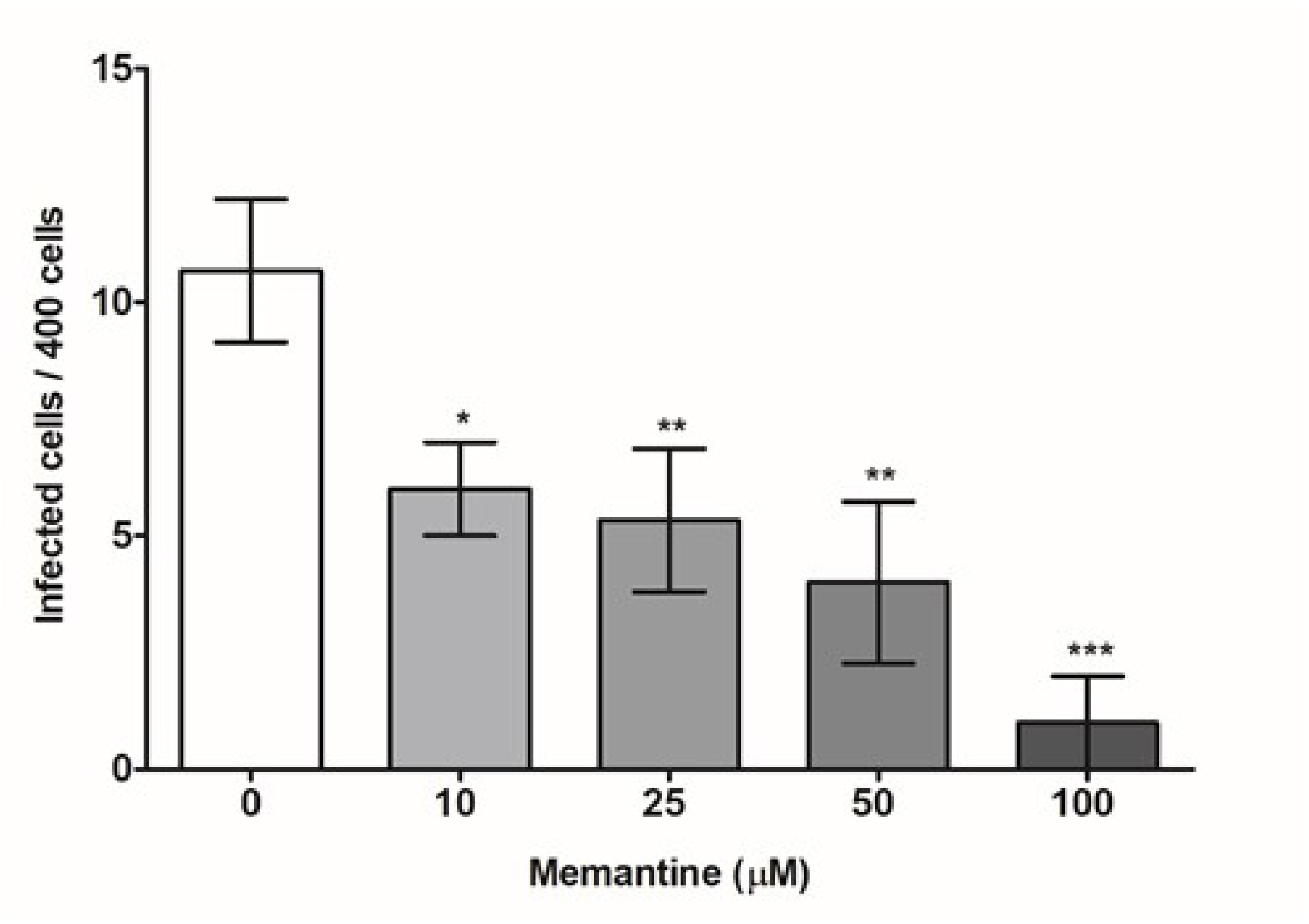
Evaluation of the effect of memantine on the intracellular cycle. RAW 264.7 macrophages (2.5×10^5^ cells/well) infected with trypomastigote forms (2.5×10^6^ parasites/well) from cell culture and incubated with different concentrations of memantine (10-100 μM) for 72 h. After this time, the cells were incubated with *Hoechst* (1: 2000) for 1 min. Cells were observed under fluorescence microscopy, using λ 350 nm excitation and λ 460 emission, and 400 cells were counted. Data are expressed as the mean ± standard deviation. * (p <0.05), ** (p <0.01). *** (p <0.001).

#### Memantine at low (but not at high) concentrations reduces NO production in RAW 264.7 macrophages *in vitro*

Due to the ability of memantine to reduce the number of infected cells in a dose-dependent manner, we were interested in checking whether memantine was acting as a trypanocidal compound by inducing macrophage activation. To evaluate the possible effect of memantine on the activation of RAW 264.7 macrophages, we first evaluated their sensitivity to LPS, a well-known macrophage activator (control). For this, the cells were incubated with different concentrations of LPS (1-100 µg/ml) for 24 h, and we considered the ability of the cells to produce NO as a measurement of activation. The NO production increased linearly with the LPS concentration; thus, among the concentrations tested, we chose 10 µg/ml (the maximum concentration tested) for further experiments (**S1A Figure**). Next, we performed a time-course experiment to follow the NO production for up to 72 h. We observed a significant increase in the period of 24 h, followed by a plateau that was maintained at 48 and 72 h (**S1B Figure**). We then evaluated the effect of memantine treatment (1-100 µM) on nitric oxide (NO) production and iNOS gene expression after 24 h of LPS stimulation. Unexpectedly, memantine treatment showed a reduction in nitrite production at concentrations of 10 and 50 μM, as well as in the expression of iNOS mRNA. Interestingly, the concentration of 100 μM did not interfere with NO production, which suggests a dose-dependent anti-inflammatory effect (**Fig 3A-B**).

**Figure 3.**
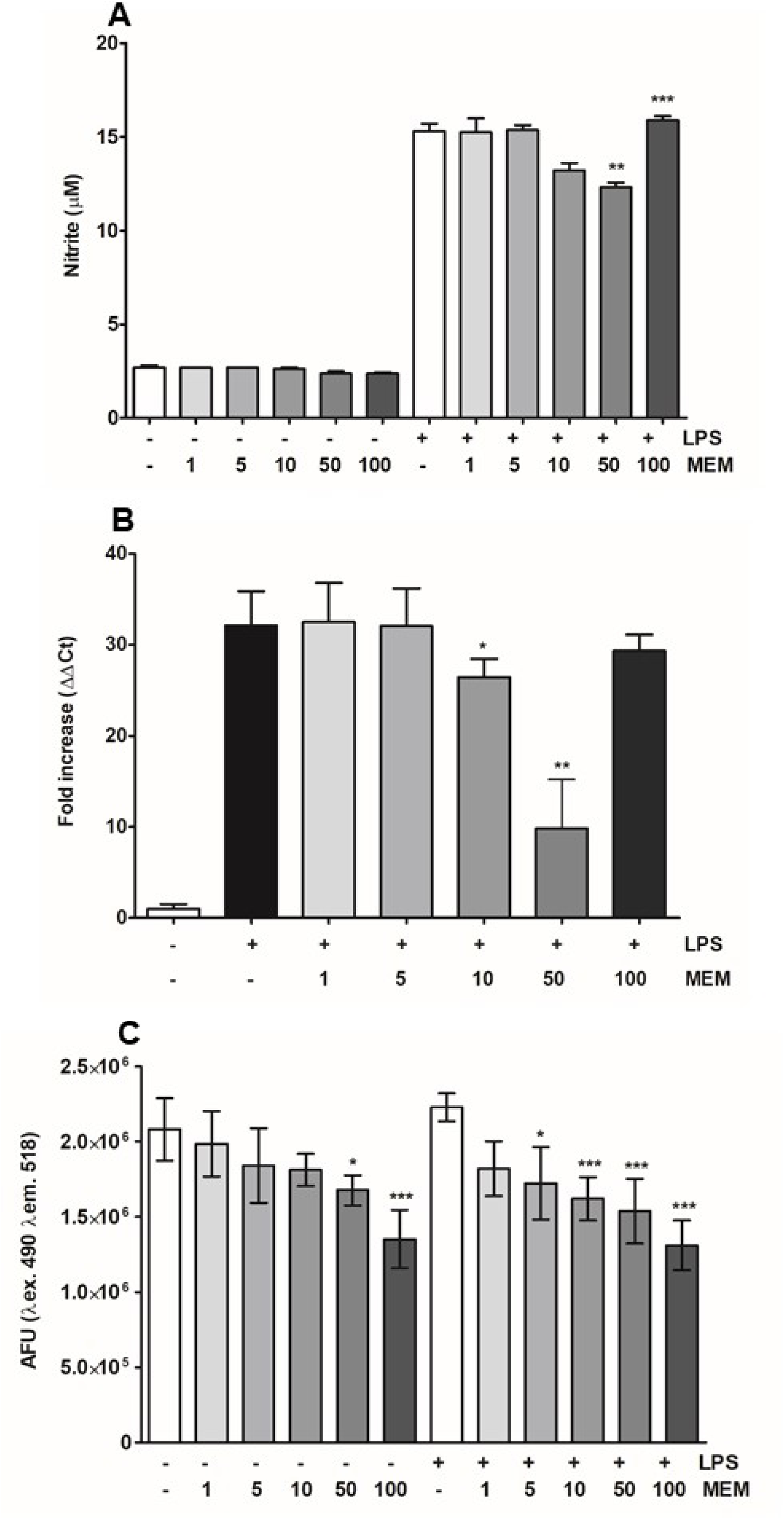
Effect of memantine on NO production and the evaluation of intracellular calcium levels. **(A)** RAW 264.7 macrophages (2.5×105 cells/well) were treated with LPS (10 μg/mL) or not treated. Cells were incubated with different concentrations of memantine (1-100 μM) for 24 h. After this period, the production of nitrites was evaluated by Griess reaction. **(B)** Gene expression of iNOS was evaluated in the presence or absence of LPS (10 µg/mL) after a 24 h incubation. Total cell RNA was extracted for cDNA synthesis and was analyzed by quantitative PCR. **(C)** RAW 264.7 macrophages (2.5×10^5^ cells/well) were treated with LPS (10 μg/mL) or not treated. Cells were incubated with different concentrations of memantine (1-100 μM) for 24 h. After this time, the cells were incubated with 5 μM fluo-4 AM for 1 h at 33 °C. The evaluation was performed on the SpectraMax i3 fluorimeter (Molecular Devices), using λ excitation 490 nm and λ emission 518. Data are expressed as a percentage ± standard deviation.* (p <0.05), ** (p <0.01), *** (p <0.001).

#### Memantine reduces intracellular Ca^2+^ levels in RAW 264.7 macrophages

It is known that the pathological activation of NMDA receptors, either by direct or indirect mechanisms, possibly results in an increase of intracellular calcium (18). Based on these observations, we evaluated the possible variations in the concentration of intracellular calcium levels in non-activated and LPS-activated RAW 264.7 cells treated with two different concentrations of memantine (1-100 μM). Exposure of the LPS-activated cells to memantine resulted in a dose-dependent decrease in intracellular calcium levels (**Fig 3C**). Remarkably, this effect was dependent on macrophage activation since no differences in the intracellular calcium levels were observed in the non-activated cells.

### In vivo

#### Memantine treatment reduces parasitemia and increases the survival of *T. cruzi*-infected BALB/c mice

Since memantine showed a reduction in the number of infected cells *in vitro*, we considered it relevant to evaluate memantine’s effect *in vivo*. The available clinical data shows the concentration administered in the treatment of patients with neurodegenerative diseases and the reported side effects for the use of memantine in other animal models (ataxia, muscle relaxation, and amnesia); the side effects were only observed in relatively high doses related to the concentration considered to be of therapeutic importance (19), and we chose 10 mg/kg of body weight as the ideal concentration for our trial. BALB/c mice infected with 1×10^3^ bloodstream trypomastigotes were treated for 10 consecutive days. The treated animals showed decreased parasitemia on the days corresponding to the parasitemic peak (7 to 10 d.p.i) by approximately 40% compared to the control group (**Fig 4A**). The mouse survival rate was 12.5% compared to the control group survival rate of 7.5% (p = 0.0347) **(Fig 4B**). In summary, memantine treatment decreased parasitemia and extended the survival of infected BALB/c mice.

**Figure 4.**
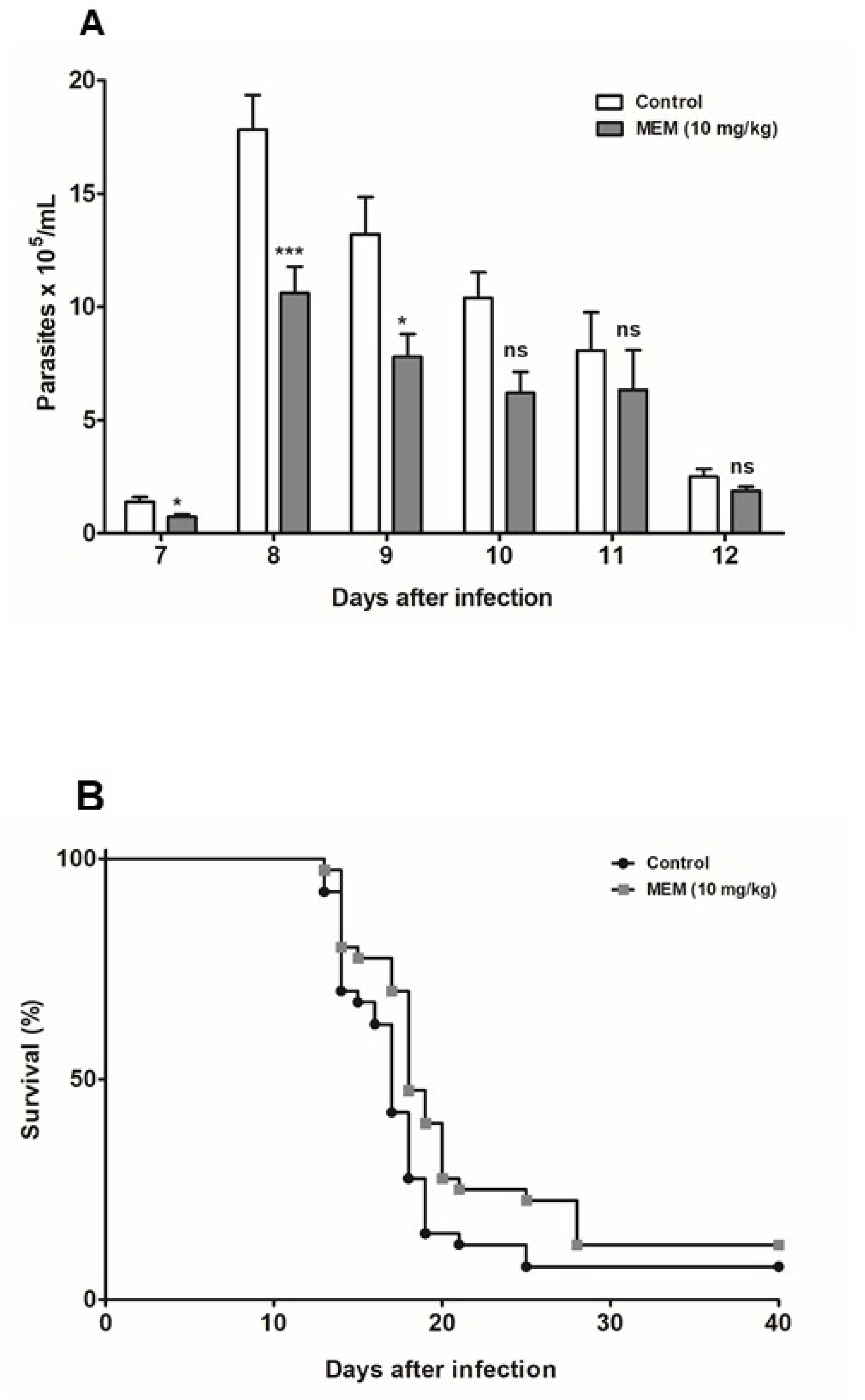
Parasitemia and mortality in treated or noninfected BALB/c mice treated with memantine (10 mg/kg per day). **(A)** Parasitemia in BALB/c mice infected by the intraperitoneal route with 1×10^3^ forms of sanguine trypomastigotes of strain Y of *T. cruzi*. The number of blood trypomastigotes on days equivalent to the parasitemic peak was evaluated in infected mice, treated or not treated with memantine (10 mg/kg per day) (N = 40). Comparison between the groups treated and not treated with memantine (p <0.05). Data are expressed as the mean ± standard deviation. **(B)** Infected animals treated or not treated with memantine (10 mg/kg per day) (N = 40). Comparison between the groups treated and not treated with memantine (log-rank test) (p <0.05).

#### The memantine treatment reduces the parasitic load and increases the inflammatory infiltrate in the heart of infected BALB/c mice

To evaluate the effect of memantine on the parasitic load in different tissues from treated mice, we performed real-time PCR to quantify the number of parasites. For this, we obtained DNA (equivalent to 5 ng of tissue DNA (P_E_/5 ng DNA) from the heart, bladder, intestine, skeletal muscle and liver. Among the evaluated tissues, the heart showed the highest parasitic load (equivalent to 3,427 ± 451 P_E_/5 ng DNA). The memantine-treated mice showed a significant reduction in the parasitic load in the heart (35.3%) compared to the parasitic load in the values obtained from the control mice (p<0.05) (**Fig 5A; S2A – E Figure**). Remarkably, the quantification of the number of amastigote nests per mm², evaluated by microscopy observation, confirmed these data: the hearts from the control mice demonstrated a mean of 2.3 ± 0.35 nests/mm^2^, while the memantine-treated mouse hearts showed a mean of 1.2 ± 0.15 nests/mm^2^ (a reduction of approximately 45%, **Fig 5B**). Moreover, we also observed that the area of the inflammatory infiltrates (normalized per mm^2^) was significantly reduced in the treated heart tissues: those from the treated animals showed a mean value of 8.66 ± 4.15 inflammatory infiltrate/mm^2^, while those from the control group presented a mean value of 80.53 ± 31.73 inflammatory infiltrate/mm^2^ (**Fig 5C**, as an illustrative example of the amastigote nests and infiltrates in treated mice vs control mice see **Fig 5D**).

**Figure 5.**
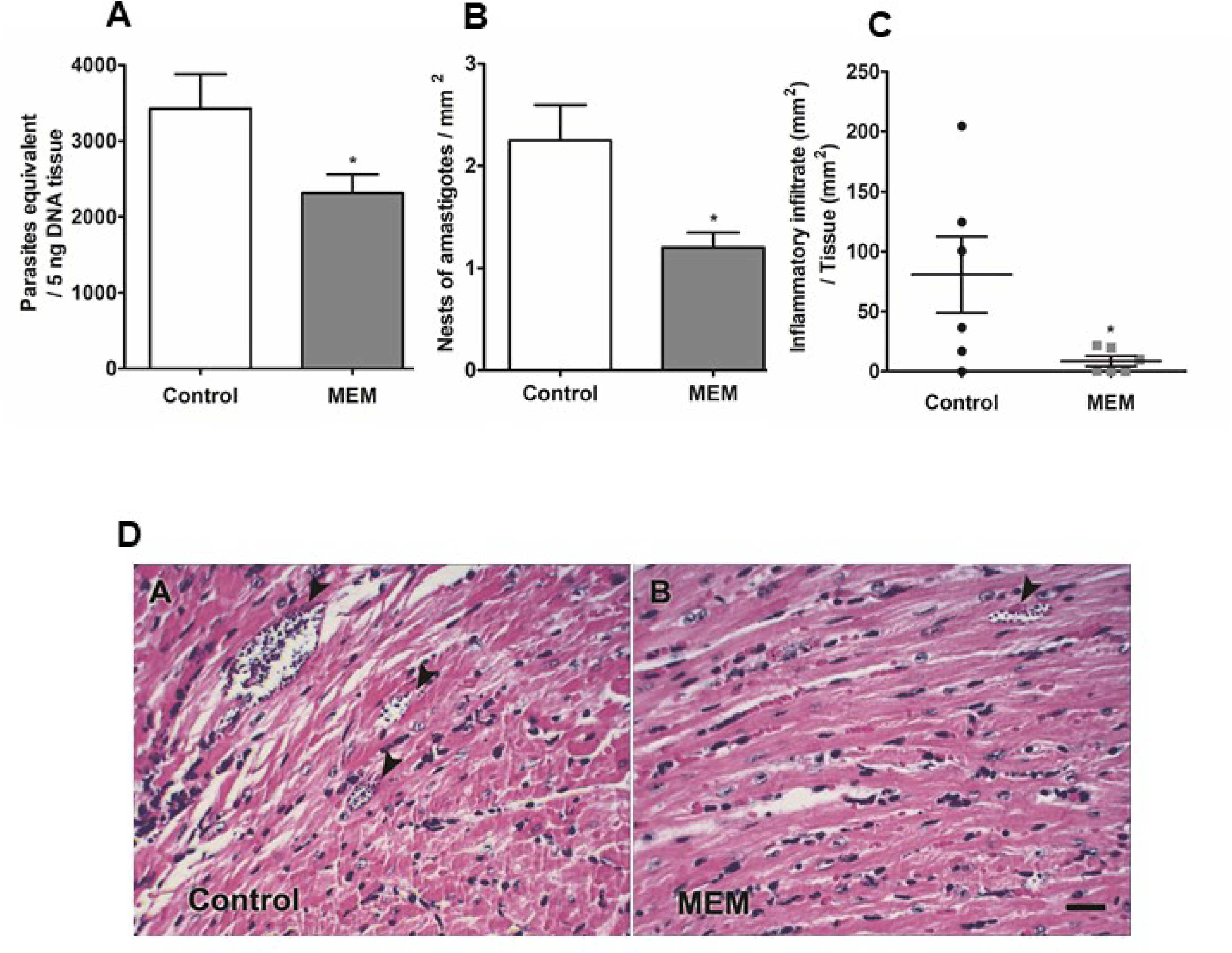
Tissue parasitic load, parasite density and inflammatory infiltrate in cardiac tissue. **(A)** Measurement of parasitic load at 15 d.p.i. in tissues of BALB/c mice infected with 1×10^3^ forms of sanguine trypomastigotes and nontreated or treated with memantine (MEM) - 10 mg/kg per day - for ten consecutive days. The graph show the number of parasites equivalent to 5 ng of tissue DNA. **(B)** Nests of amastigotes per mm^2^ in cardiac tissue at 15 d.p.i. BALB/c mice were infected with 1×10^3^ forms of blood trypomastigotes and were treated with memantine (MEM) - 10 mg/kg per day - for ten consecutive days. The data are presented in number of nests per area of the analyzed section. **(C)** Cardiac tissue sections of 5 μm thick were obtained on 15 d.p.i., stained with H & E and analyzed by light microscopy. Areas of inflammatory infiltrates were quantified by an image analysis system. The sum of infiltrated areas on the six slides was calculated for each mouse. The final individual score was expressed in square micrometers of inflammatory infiltrates per square millimeter of area examined. * (p <0.05). **(D)** Histological view of the hearts of BALB/c mice infected with sanguine trypomastigotes and nontreated or treated with memantine (MEM). The figure shows the presence of amastigotes nests (arrowheads). Scale bar represents 50 μm.

All the obtained data for the parasitic load, amastigote nest and inflammatory infiltrate quantification demonstrate that treatment with memantine reduces the risk of tissue parasitic-associated damage, both by diminishing the parasitic load and by diminishing the inflammatory response.

## Discussion

In the present work, we provide evidence of the therapeutic potential of memantine in the *in vitro* and experimental infection by *T. cruzi*. As memantine is currently used in patients with moderate to severe stages of Alzheimer’s disease (12), we propose to further study its repurposing to treat the infection by *T. cruzi*.

Under pathological conditions, memantine is used as a noncompetitive antagonist drug of the voltage-dependent N-methyl-D-aspartate (NMDA) receptor to block the effects of elevated glutamate levels (20). The NMDA receptor belongs to the family of ionotropic glutamate receptors and is involved in a variety of central nervous system (CNS) functions and processes (21). In mammals, these receptors play important physiological roles, and despite their predominance in the CNS, NMDA receptors have also been identified in peripheral and visceral sites located on the postsynaptic dendrite membranes (21, 22). In *T. cruzi*, there are no reports in the literature on the presence of a canonical NMDA-type glutamate receptor. However, as previously mentioned, our group showed that *T. cruzi* epimastigotes are responsive to NMDA (23). This information is compatible with our previous finding that the CL-14 strain of *T. cruzi* is susceptible to memantine and that amastigotes infecting CHO-K_1_ cells are the most susceptible forms *in vitro* (11). Here, we show that treatment with memantine significantly reduces the infection rate (infected/noninfected cells) in RAW 264.7 macrophages. However, as memantine diminishes NO production in infected macrophages, it is unlikely to attribute its effect of an increase in the host cell natural defense mechanism against the parasite. Therefore, other possible mechanisms altering the viability of the intracellular amastigotes were explored. Our data show that memantine induced an increase in mitochondrial function in relation to the control cells. Similarly, Prado demonstrated that Neuro-2A neural cells increased their mitochondrial reducing power when pretreated with 0.5-50 μM memantine. However, when the same cells were treated with lower memantine concentrations, the calcium influx was decreased with the concomitant increase in the mitochondrial reducing power (24). Additionally, Chen and colleagues observed that memantine at low doses may play an anti-inflammatory and neuroprotective role, although the anti-inflammatory effects are still uncertain (25). These findings corroborate our data, where memantine induced a decrease in NO production in LPS-stimulated RAW 264.7 macrophages (10 μg/mL) at concentrations of 10 and 50 μM memantine. However, the concentration of 100 μM memantine suggests a pro-oxidant effect in our assays. Additionally, memantine was shown to have potential effects as a neurotransmitter and a neuroprotective compound (26), to inhibit the ATP-sensitive potassium channels (K^+^/ATP) in substantia nigra (dopaminergic) neurons (27) and to suppress the internal currents induced by electroporation (28). In fact, Tsai and colleagues demonstrated that the concentrations used to inhibit at 50% the internal rectifying potassium channels (IK (IR)) is similar to the memantine IC_50_ (12 μM) in RAW 264.7 macrophages. These channels act as metabolic sensors and are sensitive to ATP, that is, when calcium levels are high, closure of the channel occurs (29). We previously demonstrated that memantine affects the energetic metabolism of the parasite, inducing decreased levels of ATP and triggering mechanisms that lead to apoptosis in epimastigotes of *T. cruzi* (CL strain, clone 14) (11). In the present work, we showed that 100 μM memantine induces a decrease in the intracellular calcium levels in both LPS-stimulated and nonstimulated cells. This supports the hypothesized macrophage NMDA receptor (30), suggesting that memantine would be able to block it. This is consistent with the previous observation that, in lymphocytes, NMDA receptors are involved in the regulation of intracellular calcium levels (31) as well as the levels of ROS (32).

Since memantine decreased *T. cruzi* infection in macrophages *in vitro*, we evaluated the effect of memantine treatment on infection *in vivo*. The treatment of BALB/c mice infected with Y strain bloodstream trypomastigotes with memantine for 10 consecutive days reduced the parasitemia by 40% during the parasitemic peak when compared to the control group. Importantly, the dose used in our work (10 mg/kg per day) can be considered safe: doses of 10 mg or 20 mg per day were shown to be beneficial for mice in terms of improving the functional capacity of daily activities and the behavioral disorders characteristic of Alzheimer’s disease (33).

During the acute phase of Chagas disease, it is known that the parasites are present in many tissues of the host. Our data showed a high parasitic load in the heart with 3427 ± 451 parasites equivalent to 5 ng tissue DNA (PE/5 ng DNA). As observed, memantine, at the dose used, was able to significantly reduce the tissue parasitic load in this tissue by approximately 35.3%. It was reported that animals inoculated with the trypomastigote form of the Y strain show an extremely high parasitic load on the 7th and 8th d.p.i. in the spleen and liver, among others (34). This might explain why we did not observe significant differences or a consistent parasitic load in these tissues. When we evaluated the number of amastigote nests in the cardiac tissue, our results indicate that the control group presented a mean of 2.3 ± 0.35 nests/mm^2^, while the animals treated with memantine (10 mg/kg per day) presented an average of 1.2 ± 0.15 nests/mm^2^ in cardiac tissue, which is consistent with the data obtained by using real-time PCR. Remarkably, this reduction was consistent with that observed when the inflammatory infiltrates in the cardiac tissue were analyzed. These datasets are consistent with previously published results that show the susceptibility of amastigote forms to memantine (11). This is of extreme relevance since the amastigotes are responsible for maintaining chronic infection in patients. It was shown that amastigotes can enter a dormant state, which makes them resistant to benznidazole (35). Further studies should be conducted to evaluate the possible trypanocidal effect of memantine on dormant amastigotes, which would result in an optimized alternative therapy for Chagas disease.

---

**Supporting Figure 1. Kinetics of the activation of RAW 264.7 macrophages**. **(A)** Cells of RAW 264.7 lineage macrophages (1×10^6^ cells/well) were treated with different concentrations of LPS (1-100 μg/mL). Cells were incubated for 24 hours. **(B)** Nitrite dosing in the supernatant of RAW 264.7 lineage macrophage incubated at different times (24 h, 48 h and 72 hours) with LPS (10 μg/ml). After this period, the production of nitrites was evaluated by a Griess reaction. Data are expressed as a percentage ± standard deviation. (p <0.05).

**Supporting Figure 2. Tissue parasitic load.** Evaluation of the parasitic load at 15 d.p.i. in tissues of BALB/c mice infected with 1×10^3^ forms of blood trypomastigotes and treated with memantine (MEM) - 10 mg/kg per day - for ten consecutive days**. (A)** Intestine (N = 15), **(B)** Bladder (15), **(C)** Skeletal muscle (N = 15), **(D)** Spleen (15) and **(E)** Liver (N = 15). The graphs show the number of parasites equivalent to 5 ng of tissue DNA.

